# Development and validation of the Evaluation Platform In COPD (EPIC): a population-based outcomes model of COPD for Canada

**DOI:** 10.1101/401745

**Authors:** Mohsen Sadatsafavi, Shahzad Ghanbarian, Amin Adibi, Kate Johnson, J Mark FitzGerald, William Flanagan, Stirling Bryan, Don Sin, *for the Canadian Respiratory Research Network#*

**Author notes:** Co-first authors (Mohsen Sadatsafavi, Shahzad Ghanbarian). Other collaborators: Shawn Aaron, Jean Bourbeau, Wan Tan, Teresa To, Penny Brasher, Deirdre Hennessy, Jacek Kopec. **Corresponding author:** Mohsen Sadatsafavi, Faculty of Pharmaceutical Sciences, University of British Columbia, 7th Floor, 828 West 10th Avenue, 2405 Wesbrook Mall, Vancouver, BC Canada V6T 1Z3, |. Contact for reprint request: Mohsen Sadatsafavi. This work has been presented at the Canadian Agency for Drugs and Technologies in Health 2018 Symposium, and the Society for Medical Decision Making’s 40th Annual North American Meeting.

## Abstract

**Background:** We report the development, validation, and implementation of an open-source population-based outcomes model of Chronic Obstructive Pulmonary Disease (COPD) for Canada.

**Methods:** Evaluation Platform in COPD (EPIC) is a discrete event simulation model of Canadians **40** years of age or older. Three core features of EPIC are its open-population design (incorporating projections of future population growth, aging, and smoking trends), its incorporation of heterogeneity in lung function decline and burden of exacerbations, and its modeling of the natural history of COPD from inception. Multiple original data analyses, as well as values reported in the literature, were used to populate the model. Extensive face validity as well as internal and external validity evaluations were performed.

**Results:** The model was internally validated on demographic projections, mortality rates, lung function trajectories, COPD exacerbations, and stability of COPD prevalence over time within strata of risk factors. In external validation, it moderately overestimated rate of overall exacerbations in two independent trials, but generated consistent estimates of rate of severe exacerbations and mortality.

**Limitations:** In its current version, EPIC does not consider uncertainty in the evidence. Several components such as additional (e.g., environmental and occupational) risk factors, treatment, symptoms, and comorbidity will have to be added in future iterations.

**Conclusions:** EPIC is the first multi-purpose outcome- and policy-focused model of COPD for Canada. By modeling the natural history of COPD from its inception, it is capable of modeling the outcomes of decisions across the entire care pathway of COPD. Platforms of this type have the capacity to be iteratively updated to incorporate the latest evidence and to project the outcomes of many different scenarios within a consistent framework.

## Background

Chronic Obstructive Pulmonary Disease (COPD) is one of the most common chronic diseases globally(1). The prevalence of COPD is increasing in many jurisdictions. Worldwide, the prevalence of COPD has increased by 44.2% between 1990 and 2005(1). In Canada, a country with a population of 36.7 million (as of 2017(2)), 17% of the adult population have COPD(3). COPD is characterized by the progressive and generally irreversible loss of lung function. A defining feature of COPD is the periods of intensified disease activity, referred to as acute exacerbations(4). Severe exacerbations that require inpatient care are the second leading cause of hospitalization in Canada, following only childbirth(5). COPD extolls a substantial economic burden. A Canadian study found the excess direct costs of COPD to be $5,947 (converted to 2015 Canadian dollars) per patient per year between 2001 and 2010, growing by more than 5% per year(6).

COPD is diagnosed using spirometry, and is defined as the ratio of forced expiratory volume at one second (FEV_1_) to forced vital capacity (FVC) being less than 70%, or alternatively less than the Lower Limit of Normal (LLN). The widely used Global Initiative for Chronic Obstructive Lung Disease (GOLD) severity grades are based on the ratio of FEV_1_ to its predicted value, with 80%, 50%, and 30% thresholds separating severity grades I to IV(7). These classifications are common across several other guidelines(7–9), which has resulted in a large body of evidence relating COPD severity grades to disease outcomes such as exacerbation rates and costs.

A good understanding of the future burden of diseases and the long-term consequences of clinical and policy decisions can support the search for efficiency and equity in healthcare(10). In most instances, objective projection of disease burden and outcomes are based on computer simulations(11). A common approach to predicting the future burden of diseases and the outcomes of decisions is to develop a *de novo* computer model for the specific question at hand(12). For example, different groups have created cohort-based (e.g., Markov) models of COPD to predict the future costs and outcomes of COPD in specific countries(13), to evaluate the outcomes of various strategies (e.g., improving smoking cessation rates, reducing the rate of exacerbations) for reducing the burden of COPD(14), and to evaluate the cost-effectiveness of bronchodilator therapies across different countries(15). However, this piecemeal approach has been criticized due to the resulting duplicate efforts, inconsistent assumptions, and risk of bias due to the influence of stakeholder interests when the model is developed with a particular research question in mind(16).

An alternative to this approach is to create reference models that can be used for multiple purposes(10). Reference models are the result of dedicated scientific efforts, typically with more investment integrated into the overall design, better representation of disease mechanisms and outcomes, more comprehensive evidence synthesis, and more rigorous validation exercises than the *de novo* models(17). Once developed, these reference models can be used to evaluate the outcomes of many different scenarios within a consistent framework. A number of models have been developed for COPD, and a recent systematic review has identified the strengths and weaknesses of each of the different platforms(18). The majority of these models (with some exceptions(19)) simulate a cohort of patients who initially have the disease, and therefore they cannot be used to project the future incidence of the disease and the impact of preventive scenarios such as changing smoking rates and implementation of screening programs for earlier detection of COPD.

The Evaluation Platform In COPD (EPIC) was a nationally funded research project with the aim of creating an open-source, publicly available, population-based ‘Whole Disease’ COPD model for epidemiological projections and analysis of a wide range of policies in the Canadian context. The present paper discusses the concept and overall methodology, assumptions, data sources, and results of internal and external validation.

## Methods

### Conceptual framework

The conceptual framework underlying EPIC is that of Whole Disease Modeling(16). This framework emphasizes modeling the entire pathway of the disease and the transferability of the decision node across the care pathway of the disease. It also requires the capacity for incorporating interactive decision nodes, such that implementation of a policy (e.g., the provision of a nationwide screening program for COPD) will appropriately affect the outcomes of downstream interventions (e.g., pharmacotherapy for diagnosed patients)(20).

The investigator team included key stakeholder groups in the research program including clinical experts, methodologists, and policy makers. We followed guideline recommendations on the proper conceptualization of a decision-analytic model(21). ***Figure 1*** provides a schematic illustration of the EPIC model. The following four features are the core components of the platform: 1) COPD should be modeled using the reference standard (spirometric definition) so that the model can make projections about the ‘true’ burden of COPD and the merits of screening/case detection strategies; 2) lung function trajectories and acute exacerbations are major factors impacting the natural history of COPD and should be simulated with high precision as they are ubiquitously the primary end points of clinical trials; 3) EPIC should realistically model heterogeneity in the natural history of COPD to enable modeling scenarios pertinent to Precision Medicine (e.g., the use of diagnostic and prognostic biomarkers for COPD diagnosis and treatment); 4) sex, age, and smoking, the three main risk factors for COPD, should be modeled explicitly. A consequence of the latter feature is the assumption that any changes in the prevalence of COPD over time is due to changes in the distribution of these three risk factors in the population and/or changes in the life expectancy of the population with and without COPD. Input parameters were selected from the sources that would represent ‘natural’ history of COPD, unaffected by treatment. Additional features of the disease (e.g., symptoms, treatments), other risk factors (such as environmental factors, history of asthma), and comorbid conditions will be added gradually through successive iterations.

**Figure 1:**
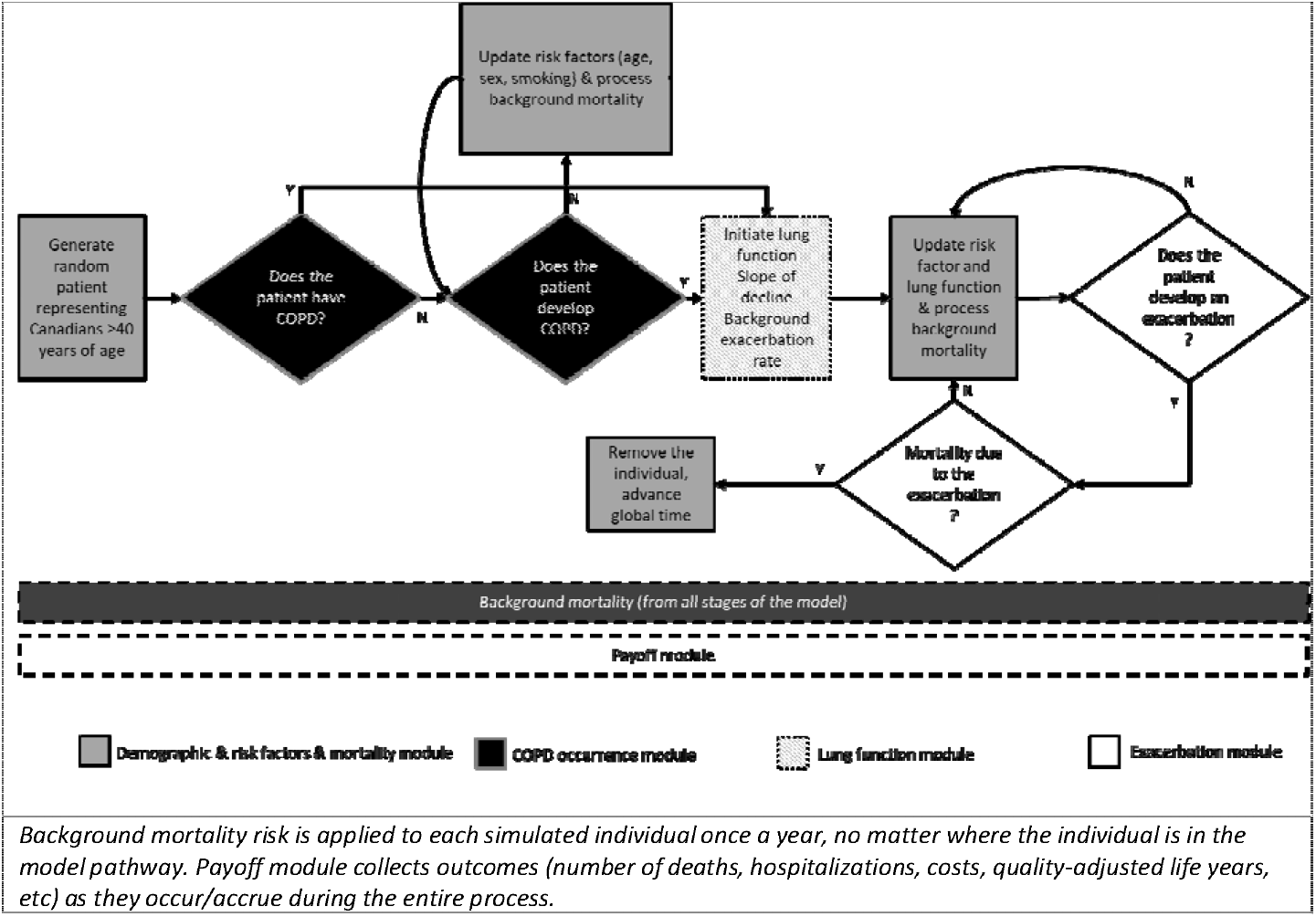
Schematic illustration of Evaluation Platform In COPD (EPIC)

EPIC is a Discrete Event Simulation (DES) of the general Canadian population. DES models the life history of individuals, one at a time, from entrance into the model to death (or end of the simulation time horizon)(22). The progression occurs during jumps in time (which is a continuous variable) from a set of pre-defined ‘events’ (e.g., COPD incidence, occurrence of an exacerbation, death – full list is available in ***Appendix 1***). An individual is defined by a set of ‘variables’ (such as sex, current age, current FEV_1_, individual-specific rate of FEV_1_ decline and rate of exacerbations) whose values determine the rate of the occurrence of each event, and events in turn affect the value of variables defining the individual. Time of events were generally sampled from the exponential distribution whose event rates were functions of an individual’s variables. Contrary to cohort-based models (such as the popular Markov models), individual-specific variables can create a virtually unlimited set of disease states, thus enabling robust modeling of heterogeneity and complex combination of disease states(22).

### Model components and sources of evidence

EPIC can be seen as a series of inter-connected components (modules), each of which models a different aspect of the disease. Current modules include 1) demographic and risk factors, 2) COPD occurrence (prevalence and incidence), 3) lung function, 4) exacerbations, 5) costs and quality of life outcomes (payoffs), and 6) mortality. ***Table 1*** provides the structure of equations for each module. The current value of parameters (i.e., regression coefficients) are provided in separate tables in the appendices.

**Table 1:**
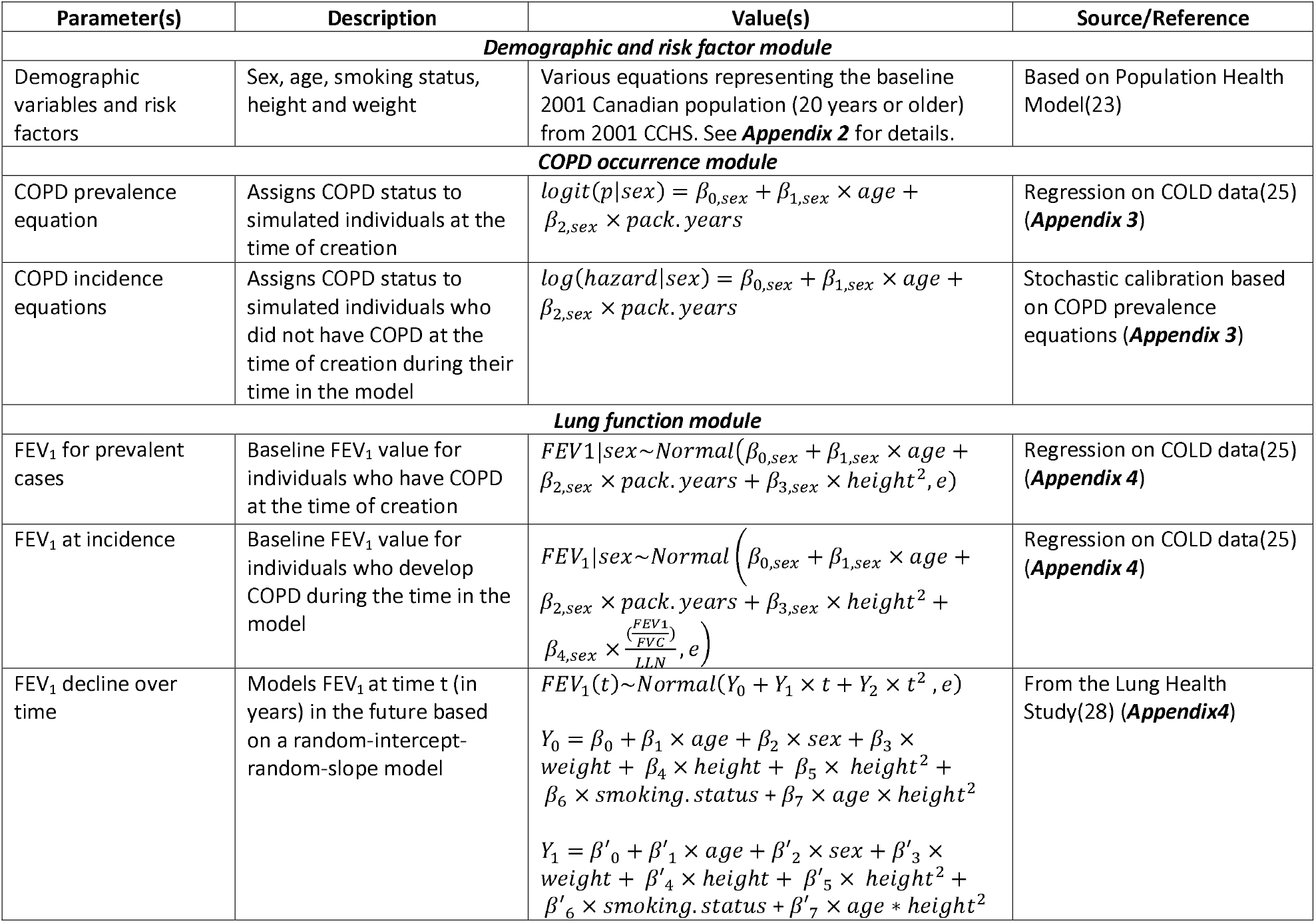
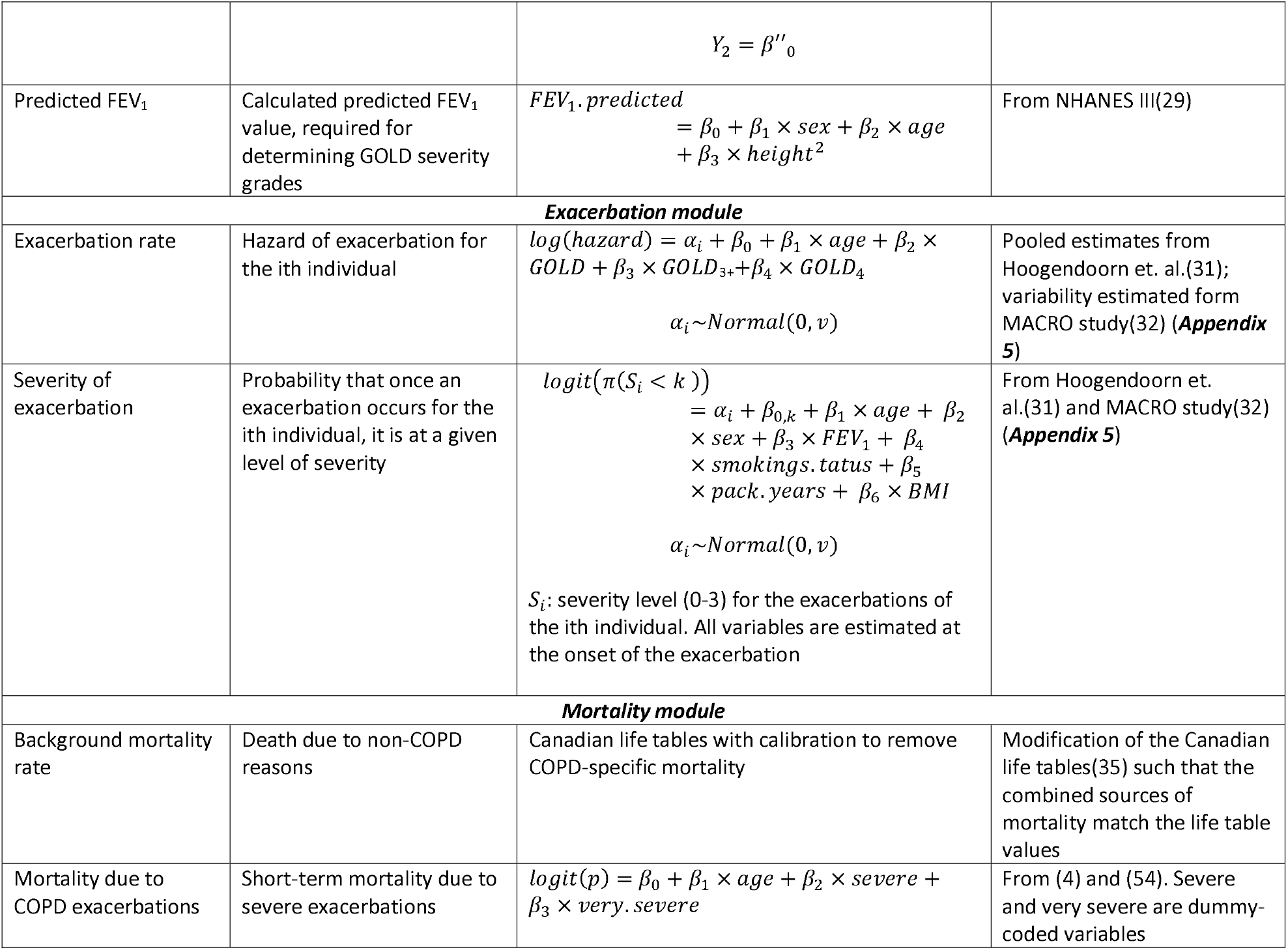
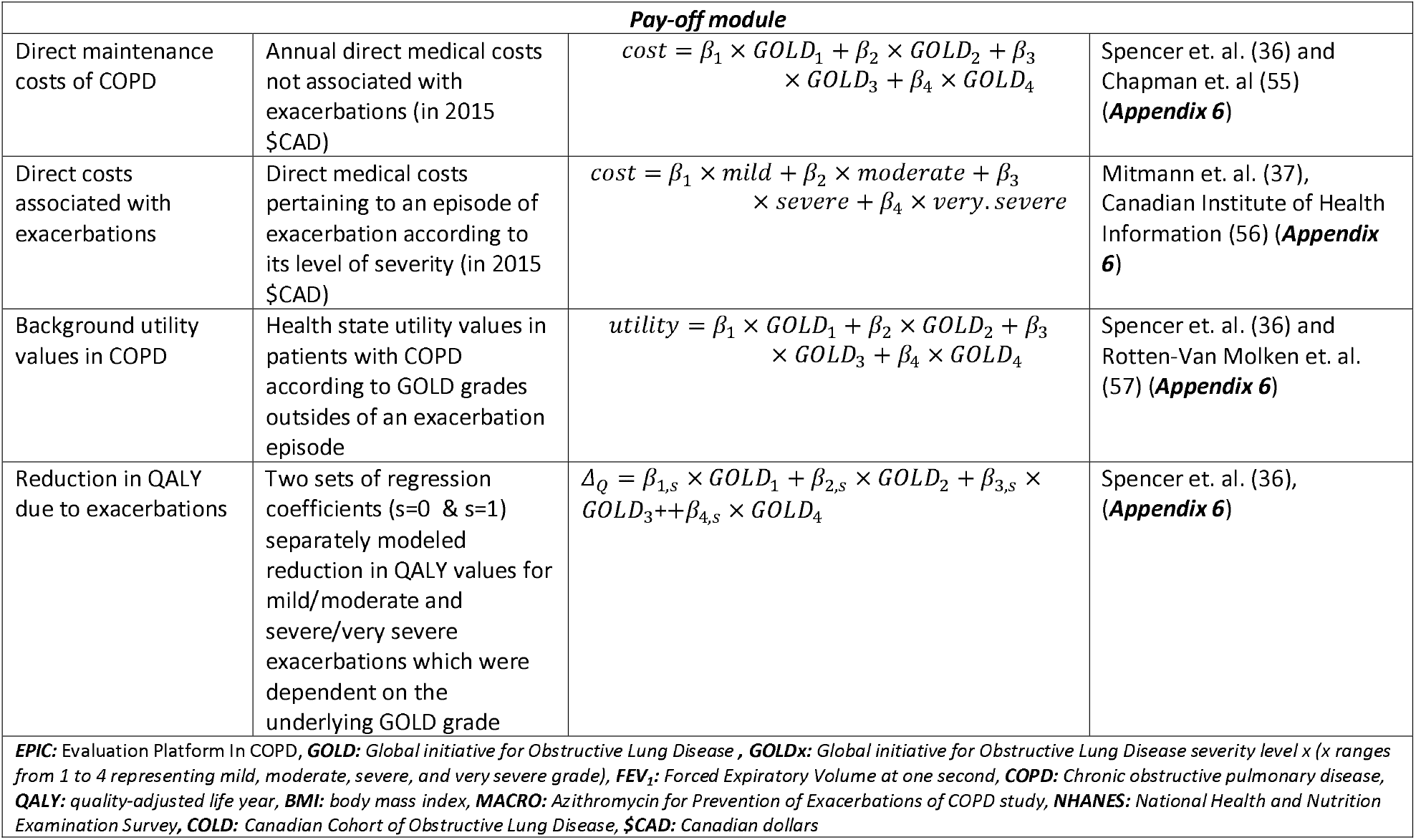
Input parameters (*β*s) and their related equations in Evaluation Platform In COPD (EPIC)

#### 1. Demographic and risk factor module

EPIC is a dynamic-population model of Canadian adults ≥40 years of age. As opposed to static models that simulate follow-up of a closed cohort of individuals for a given time (e.g., 10 years), dynamic-population models focus on the whole population within a specific date range (e.g., 2020 to 2030) and include births, deaths, immigration, and emigration. This is instrumental to the accurate modeling of temporal trends in risk factors such as population aging and changing smoking rates, and accommodating realistic aspects of policy implementations such as gradual market penetration of new medications or phased implementation of a population-based intervention such as a COPD screening program.

The demographic module of EPIC was based on Population Health Model (POHEM), a rigorously validated model of risk factors and anthropomorphic and socio-demographic characteristics (e.g., height, weight, smoking status) of Canadians, developed by Statistics Canada. POHEM has performed robustly in multiple internal and external validation challenges(23). Details of the demographic and risk factor module are provided in detail in a separate publication(23), and ***Appendix 2*** provides an overall description of the demographic module. In summary, individuals are followed over time and new individuals (incident population) enter the model in future years according to the projection of population growth and aging. This module accommodates the projected increase in life expectancy over time through the modification of future life tables. This module is also responsible for modeling smoking-related variables (current smoking status and pack-years of smoking - ***Appendix 2***). The smoking pack-years variable is dynamically updated (annually or at the time smoking status changes).

#### 2. COPD occurrence module

This module is responsible for simulating pre-existing and incident COPD. The former refers to COPD that already exists when an individual is first simulated (this is typically for the prevalent population in year 2001, and for immigrants who arrive to Canada after 2001 at older ages), while the latter occurs in simulated individuals as they age, who at the time of creation did not have COPD. COPD was defined as the ratio of FEV_1_ to FVC being below its lower limit of normal (LLN)(24). The use of LLN-based definition, as opposed to the fixed (FEV_1_/FVC< 0.70) criterion, was based on suggestions from the leading clinical experts in our team reflecting the growing support of the former definition due to its better properties (e.g., association with COPD risk factors)(24). EPIC does not model lung function in the general non-COPD population; instead, the occurrence of incident COPD is considered as an event whose risk is determined by the individual’s characteristics. Lung function is then modeled in individuals in whom the COPD switch is turned on.

##### Pre-existing COPD

We used data from the Canadian Cohort of Obstructive Lung Disease (COLD) to assign a binary COPD status to individuals upon their creation. COLD was a nationally representative, population-based cross-sectional study of 3,042 Canadians 40 years of age or older(25). Using these data, we developed sex-specific logistic regression equations for the probability of having COPD as a function of sex, age, and smoking pack-years (***Table 1*** and ***Appendix 3***).

##### Incident COPD

The sex-specific hazard of developing COPD was modeled as a log-linear function based on an individual’s sex, age, smoking pack-years, and current smoking status (***Table 1*** and ***Appendix 3***). The incident equation was derived based on a ‘steady state’ assumption: that COPD incidence is such that COPD prevalence remains stable over time within the strata of risk factors (age, sex, and smoking pack-years). These assumptions establish a mathematical relationship between incidence and prevalence(26). A stochastic optimization approach was adopted to solve for the coefficients in the incidence equation. Details of the calibration process are provided in ***Appendix 3***.

#### 3. Lung function module

Once the COPD designation is defined for an individual, an individual-specific initial FEV_1_ value and an individual-specific annual rate of FEV_1_ decline is assigned. The three components of this module are the initial FEV_1_ value for pre-existing (prevalent) COPD cases, initial FEV_1_ values for incident COPD cases, and the slope of decline in FEV_1_ over time.

For the first two components, we again relied on patient-level data from the nationally representative COLD study(27). Sex-specific linear regression equations were used to estimate FEV_1_ given risk factors (***Table 1*** and ***Appendix 4***). FEV_1_ decline was modeled as individualized equations based on the Lung Health Study (LHS, n=5,887), as described in our previous publication(28). These equations performed robustly in internal validation as well as external validation in two independent cohorts(28). In brief, we fitted a random-intercept, random-slope model for FEV_1_ decline based on 11 years of data from the LHS with covariates being age, sex, weight, height, and current smoking status (***Table 1*** - given that the original equation had other risk variables such bronchial hyper-responsivity not modeled in EPIC, we refitted the original model with reduced variables – results are provided in ***Appendix 4***). An additional quadratic term for time was included to capture the potential acceleration in lung function decline over time.

##### GOLD severity grades

Much of the evidence on COPD outcomes (e.g., costs, quality of life, exacerbation rates) is available according to the GOLD severity grades which are based on FEV_1_ to its predicted ratio. Calculating the predicted FEV_1_ is based on individuals’ age, sex, and height using the National Health and Nutrition Examination Survey equations(29).

#### 4. Exacerbation module

The exacerbation module simulates the exact timing of acute exacerbations of COPD, as well as their severity. Exacerbations were defined in four severity levels according to the event-based definition of exacerbations (which is based on healthcare services use)(30): mild exacerbations are those that result in the intensification of symptoms but do not require an encounter with the healthcare system; moderate exacerbations are those in which patient visits a physician or emergency department but is not hospitalized; severe exacerbations are those that result in hospital admissions; and very severe exacerbations are those that result in admission to the Intensive Care Unit.

We used separate sources for estimating the average rate of exacerbations and between-individual variability (heterogeneity) in rates (***Table 1***, ***Appendix 5***). The average rate was modeled as a function of the GOLD severity grades according to the pooled analysis by Hoogendoorn et. al.(31). The ratio of severe/very severe to total exacerbations was selected from the same study. However, this source did not provide the ratio of mild to moderate or severe to very severe exacerbations. These were selected from a dedicated analysis of a large one-year clinical trial of azithromycin therapy in preventing exacerbations (The MACRO study, n=1,142)(32). The instantaneous rate of exacerbation (hazard) was modeled as a log-linear function of variables; similarly, a proportional odds model was used to assign probabilities for various exacerbation severity levels (***Table 1***).

In EPIC, simulated individuals have their specific rate of exacerbations, as well as their specific probability for an exacerbation, once it occurs, to be at a given level of severity. Empirical evidence supports the presence of such heterogeneity(33). The extent of such heterogeneity was based on our previous work using data from the MACRO study(34). This heterogeneity was modeled through individual-specific (random-effect) intercepts for log-hazard of exacerbation rate and log-odds of exacerbation severity (***Table 1***)(34).

#### 5. Mortality module

This module models death from two sources: due to severe or very severe COPD exacerbations, and background mortality (due to other causes). Background mortality rates were taken from Canadian Life Tables and incorporated the projected declining mortality rates in the future(35). In addition, the probability of death from severe or very severe exacerbations was obtained from the literature (***Table 1***). Explicit modeling of COPD-related mortality in EPIC required calibration of background mortality rates so that the combined sources of mortality would produce mortality rates that resemble the sex-specific Canadian life tables. Finally, to avoid implausibly low FEV_1_ values, and based on the opinion of the clinical leaders of the team, we assumed individuals would die of a severe exacerbation once they reach an FEV_1_ of 0.5.

#### 6. Payoff module

The payoff module assigns costs and health state utility values to events and collects the aggregate population values. These values were chosen from the literature after a dedicated literature review. Details of costs and utility calculations are provided in ***Appendix 6***. EPIC currently does not model medical costs and quality of life in the general non-COPD population. Only COPD-specific direct medical costs and utilities are considered. Two cost components are modeled. Annual ‘maintenance’ costs of COPD are those that accrue outside of episodes of exacerbations. These costs were modeled according to the GOLD grades from previously published studies(36). The second component is costs associated with exacerbations, which were modeled separately for each level of exacerbation severity based on Canadian studies(14,37). All costs are adjusted to 2015 Canadian dollars using the healthcare component of the Consumer Price Index. Similarly, two sets of utilities were modeled: utilities outside exacerbation episodes according to GOLD grades from the published literature, and a direct reduction in QALY associated with exacerbations given their level of severity(36).

### Simulation platform

EPIC was programmed in C++. Two copies of the model were developed: one in Statistics Canada’s Model Generator, a set of libraries and header files for DES programming implemented in Microsoft^®^ Visual Studio, with pre-existing code representing POHEM(38). An identical open-source and open-access model was developed from scratch with an interface in R (v3.5.1)(39). EPIC code is publicly available (at http://resp.core.ubc.ca/research/Specific_Projects/EPIC) and can be downloaded and run as an R package.

### Examining face validity and internal validity

We followed the guidelines in establishing validation targets(40). Face validity refers to whether the model produces sensible results and if changing input parameters affects model output in the expected direction. Two important sets of face validity targets were the gradient of outcomes across categories of risk factors (e.g., higher COPD prevalence among ever-smokers than never-smokers), and the stability of outcomes over time within risk factor strata. Face validity results are not reported here but are available upon request. Internal validity was assessed by examining if the model produced results that matched the patterns observed in input data sources or calibration targets. These included **1)** comparing the simulated population growth and aging with Statistics Canada’s projections (validating demographic and mortality modules), **2)** comparing age-specific all-cause mortality rates with Statistic Canada’s life table estimates (validating the mortality module), **3)** comparing COPD prevalence and GOLD grades by risk factor groups with estimates from the COLD study (validating the COPD occurrence and lung function modules), **4)** comparing average and 95% prediction interval of FEV_1_ decline with prediction equations from LHS (validating the lung function module), **5)** comparing the rate of total and severe exacerbations per GOLD severity grades with the original pooled estimates by Hoogendoorn et. al. (31) and results from the MACRO study (validating the exacerbation module), and **6)** comparing the average costs and QALYs across GOLD grades with expected values that are affected by background values and changes due to the occurrence of exacerbations (validating the pay-off module).

### Examining external validity

Ensuring external validity requires examining the model output against results from external sources of evidence that were not used in the model as input parameters or calibration targets. The external validity tests were based on the reported outcomes from the placebo arms of two independent clinical trials: TOwards a Revolution in COPD Health (TORCH)(41) and Understanding Potential Long-Term Impacts on Function with Tiotropium (UPLIFT)(42). These studies have been used by other COPD modeling groups as external validity targets and their use here also provides cross-validity tests between EPIC and existing COPD models(18,19,43–47). The external validity tests involved creating samples that mimicked the study sample of TORCH and UPLIFT to compare three quantities: rate of overall exacerbations, rate of severe exacerbations, and percentage of patients who died during follow-up. Rejection sampling was used to create EPIC samples that matched the baseline characteristics (age, sex, smoking status, FEV_1_, and GOLD grades) of patients in TORCH and UPLIFT (details in ***Appendix 7***).

## Results

### Model calibration and internal validation

#### Demographic and mortality modules

After incorporating COPD-specific mortality and calibrating background mortality, the model generated age-specific all-cause mortality rates that were close to the Canadian life tables (***Appendix 2 - Figure 2.1***). The model performed robustly in predicting population growth and aging in the future. ***Appendix 2 - Figure 2.2*** presents the simulated (EPIC’s) and projected (Statistics Canada’s) population growth, as well as population age pyramid for three exemplary years (2015, 2025, and 2034,).

#### COPD occurrence module

After the model was calibrated using the methodology presented in ***Appendix 3***, the coefficient for calendar year in the logistic model of COPD prevalence (controlling for sex, age, and smoking) was - 0.0004 for males and 0.0001 for females, indicating that the incidence and prevalence equations were coherent. The model performed robustly in replicating the COPD prevalence by sex and age groups as observed in the COLD study (***Figure 2***).

**Figure 2:**
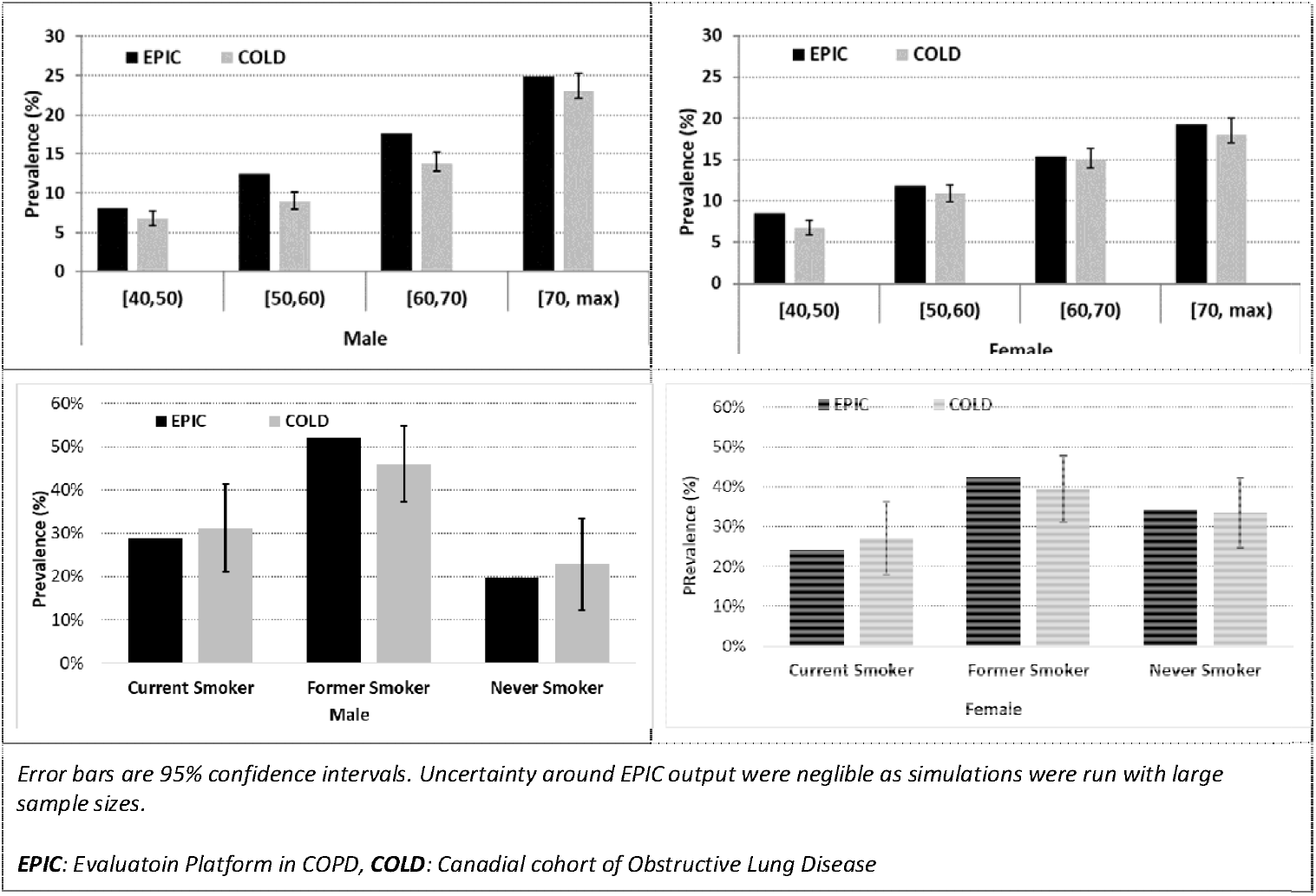
Comparison of the results of EPIC for COPD prevalence by sex and age groups with estimations from COLD data (the source of prevalence equations)

#### Lung function module

The equation for FEV_1_ decline had shown robust internal validity (against LHS) as well as external validity against two independent cohorts. Details of the validation process for FEV_1_ equations are provided in its original report(28). Given these results, to validate this module we compared the simulated FEV_1S_ from EPIC against the LHS-based equations. ***Figure 3*** presents the results across four subgroups of subjects (based on permutations of sex and smoking status at baseline). Overall, the FEV_1_ values simulated in EPIC were similar to the LHS-based predictions in terms of the mean and standard deviation across the range of possible values. The probabilities that the FEV_1_ value simulated in EPIC fell within the LHS-based 95% prediction interval were between 94% and 96% for all four subgroups.

**Figure 3:**
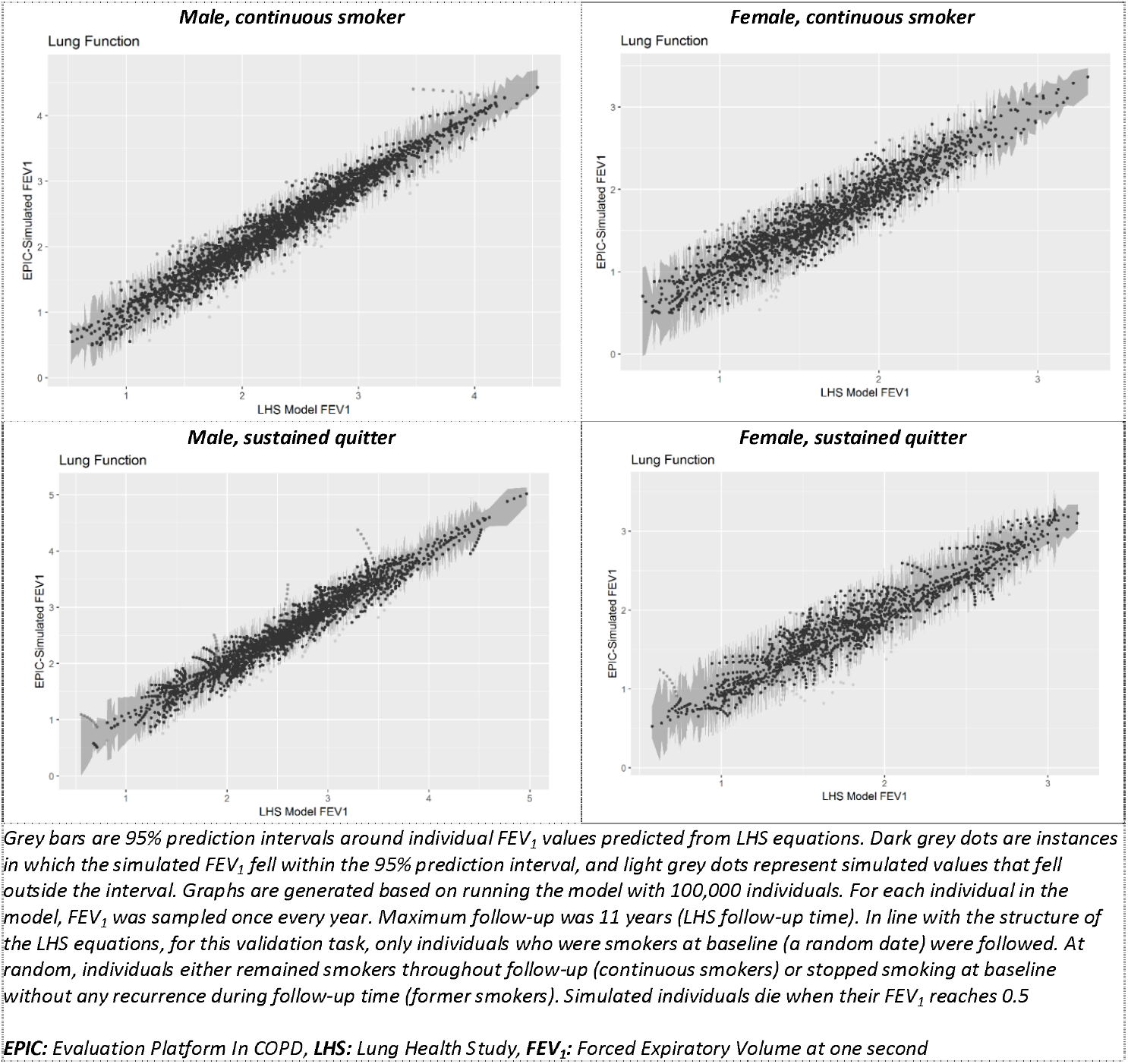
Scatterplot of simulated FEV_1S_ from EPIC versus predicted FEV_1S_ from LHS equations for up to 11 years

#### Exacerbation module

***Figure 4*** shows the EPIC-simulated annual rate of total and severe exacerbations stratified by GOLD grade compared with those reported in Hoogendoorn et. al. (31). EPIC mildly underestimated the rate of both total and severe exacerbations in GOLD grade I; nonetheless the simulated values were within the 95% confidence intervals (95%CI) of the original pooled estimates.

**Figure 4:**
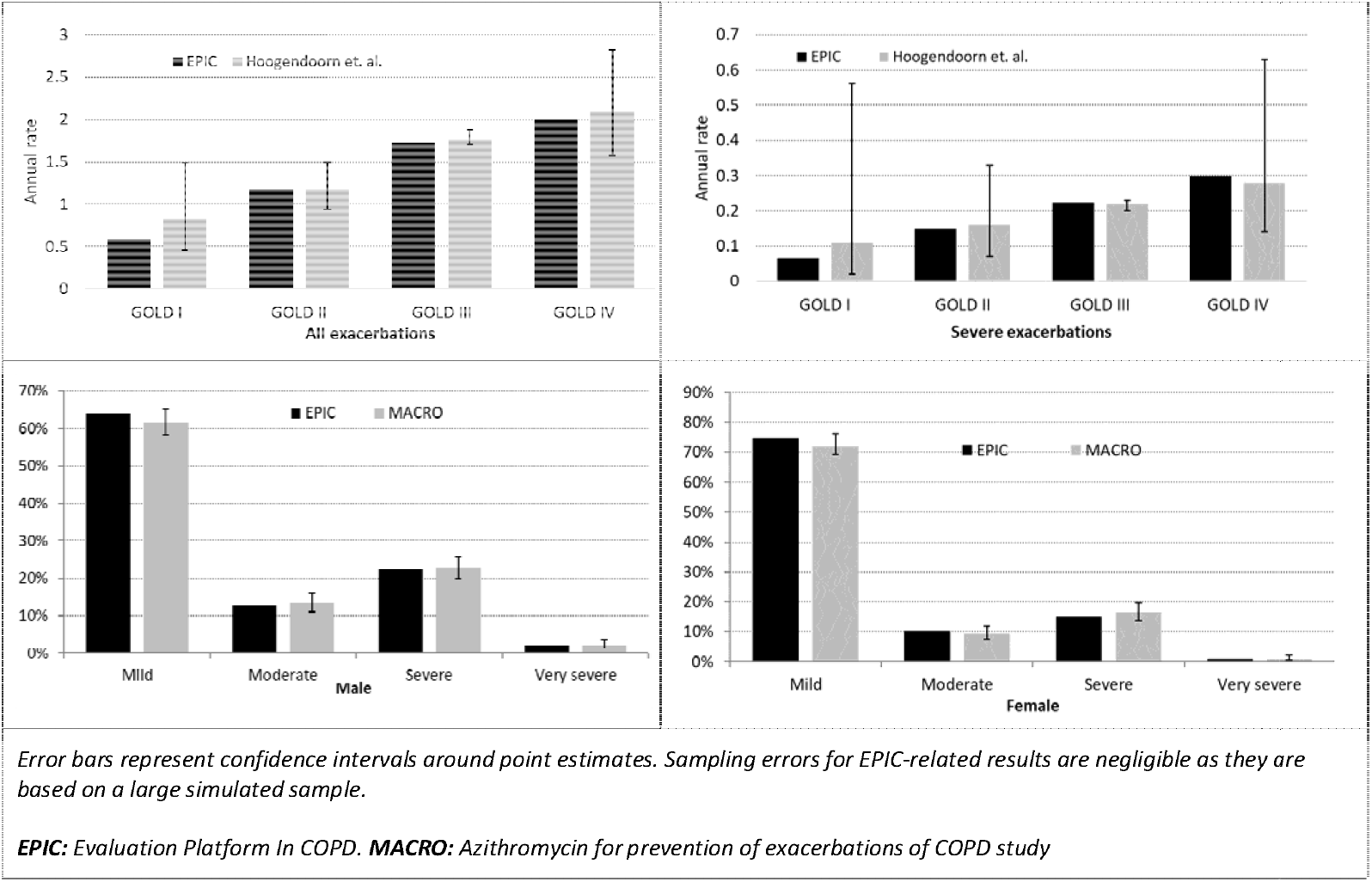
Top panel: Comparison of the results of EPIC for the rate of all (left) and severe (right) exacerbations with the pooled estimates from Hoogendoorn et. al. Bottom panel: comparison between the distribution of exacerbation severity between EPIC and MACRO study for males (left) and females (right)

#### Pay-off module

***Appendix 6 – Figure 6.1*** provides the results of internal validation of the pay-off module. Back-calculating costs and QALYs given their background values and the expected rate of exacerbations (of each level of severity) generated estimates that were close to the output of EPIC.

### External validation

After rejection sampling, the baseline characteristics of the simulated sample in EPIC resembled those of the original samples of TORCH and UPLIFT (***Appendix 7***). ***Figure 5*** provides the results of external validation tests.

**Figure 5:**
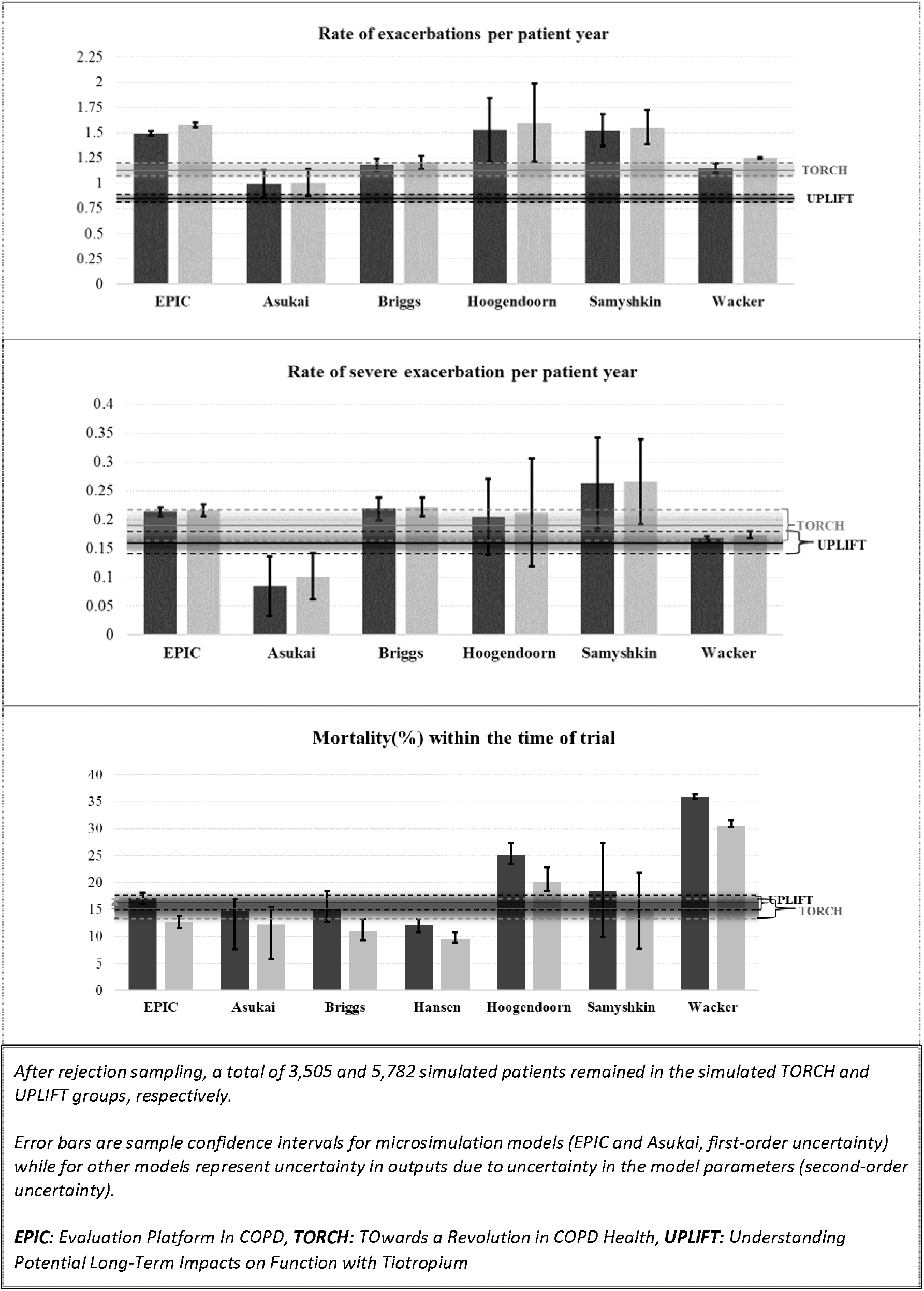
Comparison of the results of EPIC and five different COPD models with TORCH (light grey bars) and UPLIFFT (dark grey bars) for total exacerbations (top panel), severe exacerbation rate (middle panel), and mortality rate (bottom panel)

EPIC, along with two other COPD models (19,47) produced a higher total exacerbation rate than was observed in the UPLIFT trial (1.55/patient-year in EPIC versus 0.85/patient-year in UPLIFT, top panel of ***Figure 5***). Similarly, EPIC and the same two models overestimated exacerbation rate in TORCH (1.51/patient-year in EPIC versus 1.13/patient-year in TORCH). EPIC-generated values were outside of the 95%CI of both studies. However, EPIC did replicate the higher exacerbation rate observed in TORCH compared with UPLIFT. EPIC simulated a slightly higher rate of severe exacerbations than was observed in TORCH and UPLIFT (0.21/patient-year in EPIC versus 0.17/patient-year in UPLIFT, and 0.22/patient-year in EPIC versus 0.19/patient-year in TORCH, middle panel of ***Figure 5***). EPIC-generated estimate was outside of the 95%CI of UPLIFT. Three other models (19,45,47) generated higher rates than was observed in the trials. EPIC simulated a lower mortality rate over the trial follow-up (12.8%, bottom panel in ***Figure 5***) than was observed in TORCH (15.2%); EPIC-generated value was outside of the 95%CI. EPIC generated a similar mortality rate (17.3%) to what was observed in UPLIFT (16.3%).

## Discussion

We reported the conceptualization, development, implementation, as well as internal and external validation of a population-based microsimulation outcomes model of COPD in Canada. This platform was developed to provide the capacity to project the burden of COPD given changing risk factors and arrival of new technologies for primary, secondary, and tertiary care prevention. The major components of the model were simulated life trajectories of Canadians, equations describing the incidence and prevalence of spirometrically-defined COPD, the individual-specific progression of lung function decline in COPD, the individual-specific rate and severity of COPD exacerbations, COPD-related mortality rates, and estimates of costs and quality of life weights associated with different levels of disease severity and exacerbation status. The use of an open population is a key feature of the model as it provides the capacity to project future outcomes while considering temporal trends in risk factors such as aging and smoking. It also enables gradual market penetration of interventions and adoption of policies to be modeled. This was made possible by access to the population-based cohort in which COPD was defined spirometrically (as opposed to clinical cohorts that are typically based on convenience sampling of diagnosed patients in the community). Modeling heterogeneity in key components of the disease mechanisms (lung function decline and exacerbations) is another important feature of EPIC which should facilitate the evaluation of scenarios pertaining to complex decision rules for Precision Medicine. In developing EPIC, we placed a major emphasis on using the most relevant sources of evidence for different components of the model. This included an extensively validated pre-existing model of sociodemographic characteristics of the Canadian population and trends in the distribution of risk factors (e.g., smoking), a secondary analysis of the landmark LHS study for modeling FEV_1_ trajectories, a secondary analysis of a population-based Canadian study for modeling COPD incidence and prevalence, and a secondary analysis of a recent clinical trial to quantify heterogeneity in the burden of exacerbations.

Overall, the model performed robustly in face validity and internal validity tests. This should be reassuring in terms of the validity of the original statistical analyses, model calibration exercises, and model implementation. It also performed satisfactorily in the external validity of the against two independent studies that have also been used to validate other COPD models. It overestimated the rate of overall exacerbation, but its simulated exacerbation rates were similar to two other COPD models. For rate of severe exacerbations and proportion of patients who died during the follow-up period of original studies, its predictions were better aligned with the reported values and were generally around the mid-point of simulated values from five other COPD models. The model replicated the direction of the differences in exacerbation and mortality rates between the two trials, which indicates its responsiveness in propagating changes in risk factors to changes in outcomes.

The limitations of EPIC also need to be mentioned. This work represents the core structure of a COPD ‘Whole Disease’ model, but much work needs to be done to create a comprehensive COPD outcomes model. While in its current form EPIC is capable of making projections about the future burden of COPD for a broad range of policies, there are many other aspects of COPD which need to be incorporated. Examples include treatment, symptoms, diagnosis, and comorbid conditions. Further, EPIC does not currently incorporate uncertainty in the underlying evidence, which is a goal for the next iteration of this platform. Given that many parameters of EPIC are derived based on calibration techniques, this is not simply a matter of replacing fixed input parameters with their probabilistic equivalent. More sophisticated model calibration techniques might prove useful in this context(48). Further, EPIC is currently limited to modeling the impact of sex, age, and smoking as risk factors for COPD. As such, it cannot currently be used to evaluate scenarios that relate to occupational and environmental exposures or history of asthma. Because lung function in the non-COPD population was not modeled, preventive strategies based on monitoring lung function prior to persistent airflow obstruction cannot be assessed. Similarly, prodromal phase of exacerbations and exacerbation phenotypes (e.g., infectious versus inflammatory) are not currently modeled, limiting the applicability of EPIC in simulating scenarios pertaining to the early detection of exacerbations. Overall, EPIC can currently be used for modeling the outcomes of disease management based on lung function, exacerbation history, and smoking history. However, it cannot provide this capacity for disease management based on symptoms or burden of comorbidity. The planned iterative upgrades should see these features gradually incorporated into the platform.

EPIC is not the first multi-purpose model of COPD. We have previously performed a systematic review of published COPD models that were used for epidemiological projections or economic evaluations(49). The majority of models were created for addressing a specific question (e.g., cost-effectiveness of a novel therapy). However, in some instances, subsequent models were built on previous ones in an iterative fashion. The presence of such an identifiable ‘lineage’ of models demonstrates the need for, and feasibility of, re-using an existing model for different objectives. Recently, a convention of model developers assessed the consistency of five different models in COPD and their external validity(18,50,51). Putting our developed model in the context of previous reference models is informative. Among the previously reviewed models, only the model by Hoogendoorn et. al., which was developed for the Netherlands, is a dynamic population model(19). EPIC appears to be unique in modeling heterogeneity at the individual level in lung function and the rate and severity of exacerbations. Briggs et. al., developed a model of COPD progression based on the Evaluation of COPD Longitudinally to Identify Predictive Surrogate End-points (ECLIPSE)(52) data (with resource-use estimates from UPLIFT)(45). The model was based on a rigorous practice of model conceptualization(53), and has performed satisfactorily in internal and external validation studies(18,45). A major advantage of this model is its incorporation of multiple aspects of COPD (such as symptoms and six-minute walking distance, a measure of functional capacity). However, reliance on mainly one study (ECLIPSE) makes this model less relevant to the Canadian context, and three-year follow-up of ECLIPSE makes long-term projections less certain.

Predicting the future using the best available evidence is a cornerstone of many disciplines. In healthcare, the use of modeling to reconcile evidence from multiple sources to project the outcomes of decisions is considered “an unavoidable fact of life”(11). Model-based projections can critically inform recommendations for best practices as well as policies. Multi-purpose models that encapsulate concerted efforts in evidence synthesis, validation, and implementation exist for many health conditions. EPIC adds to the portfolio of such COPD models currently under development by various groups(51). While EPIC is developed for the Canadian context, many of its components pertain to the natural history and biology of COPD and are independent of a particular setting. Other components can be updated (mainly by changing numerical values of input parameters rather than modifying the structure of equations) with the specifics of a healthcare setting. As such, this platform can have applicability for other jurisdictions. The recent organized efforts in ensuring external validity and cross-comparability of such models can help the development of robust platforms that address the ongoing needs of both the research and policy community.

## References

1. Soriano JB, Abajobir AA, Abate KH, Abera SF, Agrawal A, Ahmed MB, et al. Global, regional, and national deaths, prevalence, disability-adjusted life years, and years lived with disability for chronic obstructive pulmonary disease and asthma, 1990–2015: a systematic analysis for the Global Burden of Disease Study 2015. Lancet Respir Med [Internet]. 2017 Aug [cited 2017 Aug 22]; Available from: http://linkinghub.elsevier.com/retrieve/pii/S221326001730293X

2. Statistics Canada. Canada at a Glance - 2018 - Population [Internet]. [cited 2018 Oct 1]. Available from: https://www150.statcan.gc.ca/n1/pub/12-581-x/2018000/pop-eng.htm

3. Evans J, Chen Y, Camp PG, Bowie DM, McRae L. Estimating the prevalence of COPD in Canada: Reported diagnosis versus measured airflow obstruction. Health Rep. 2014 Mar;25(3):3–11.

4. Aaron SD. Management and prevention of exacerbations of COPD. BMJ. 2014;349:g5237.

5. Camp P, Levy R. A snapshot of chronic obstructive pulmonary disease in British Columbia and Canada. BCMJ. 2008;50(2):80.

6. Khakban A, Sin DD, FitzGerald JM, Ng R, Zafari Z, McManus B, et al. Ten-Year Trends in Direct Costs of COPD: A Population Based Study. Chest. 2015 Jun 4;148(3):650–646.

7. From the Global Strategy for the Diagnosis, Management and Prevention of COPD. Global Initiative for Chronic Obstructive Lung Disease (GOLD) 2017 [Internet]. Available from: http://goldcopd.org

8. American Thoracic Society/European Respiratory Society Task Force. Standards for the Diagnosis and Management of Patients with COPD [Internet]. [Internet]. New York: American Thoracic Society; [cited 2014 Nov 27]. Report No.: Version 1.2. Available from: http://www.thoracic.org/clinical/copd-guidelines/

9. Gruffydd-Jones K, Loveridge C. The 2010 NICE COPD Guidelines: how do they compare with the GOLD guidelines? Prim Care Respir J J Gen Pract Airw Group. 2011 Jun;20(2):199–204.

10. Scotland G, Bryan S. Why Do Health Economists Promote Technology Adoption Rather Than the Search for Efficiency? A Proposal for a Change in Our Approach to Economic Evaluation in Health Care. Med Decis Mak Int J Soc Med Decis Mak. 2017;37(2):139–47.

11. Buxton MJ, Drummond MF, Van Hout BA, Prince RL, Sheldon TA, Szucs T, et al. Modelling in economic evaluation: an unavoidable fact of life. Health Econ. 1997;6(3):217–27.

12. Bellete B, Coberly J, Barnes GL, Ko C, Chaisson RE, Comstock GW, et al. Evaluation of a whole-blood interferon-gamma release assay for the detection of Mycobacterium tuberculosis infection in 2 study populations. Clin Infect Dis Off Publ Infect Dis Soc Am. 2002 Jun;34:1449–1456.

13. Nielsen R, Johannessen A, Benediktsdottir B, Gislason T, Buist AS, Gulsvik A, et al. Present and future costs of COPD in Iceland and Norway: results from the BOLD study. Eur Respir J Off J Eur Soc Clin Respir Physiol. 2009 Oct;34(4):850–7.

14. Najafzadeh M, Marra CA, Lynd LD, Sadatsafavi M, FitzGerald JM, McManus B, et al. Future Impact of Various Interventions on the Burden of COPD in Canada: A Dynamic Population Model. PLoS ONE. 2012 Oct 11;7(10):e46746.

15. Oostenbrink JB, Rutten-van Mölken MPMH, Monz BU, FitzGerald JM. Probabilistic Markov model to assess the cost-effectiveness of bronchodilator therapy in COPD patients in different countries. Value Health J Int Soc Pharmacoeconomics Outcomes Res. 8:32–46.

16. Tappenden P, Chilcott J, Brennan A, Squires H, Stevenson M. Whole disease modeling to inform resource allocation decisions in cancer: a methodological framework. Value Health. 2012 Dec;15(8):1127–36.

17. Afzali HHA, Karnon J, Merlin T. Improving the accuracy and comparability of model-based economic evaluations of health technologies for reimbursement decisions: a methodological framework for the development of reference models. Med Decis Mak Int J Soc Med Decis Mak. 2013 Apr;33(3):325–32.

18. Hoogendoorn M, Feenstra TL, Asukai Y, Borg S, Hansen RN, Jansson S-A, et al. Cost-effectiveness models for chronic obstructive pulmonary disease: cross-model comparison of hypothetical treatment scenarios. Value Health J Int Soc Pharmacoeconomics Outcomes Res. 2014 Jul;17(5):525–36.

19. Hoogendoorn M, Rutten-van Mölken MPMH, Hoogenveen RT, van Genugten MLL, Buist AS, Wouters EFM, et al. A dynamic population model of disease progression in COPD. Eur Respir J Off J Eur Soc Clin Respir Physiol. 2005 Aug;26(2):223–33.

20. Birch S, Gafni A. Cost effectiveness/utility analyses. Do current decision rules lead us to where we want to be? J Health Econ. 1992 Oct;11(3):279–96.

21. Roberts M, Russell LB, Paltiel AD, Chambers M, McEwan P, Krahn M, et al. Conceptualizing a model: a report of the ISPOR-SMDM Modeling Good Research Practices Task Force--2. Value Health J Int Soc Pharmacoeconomics Outcomes Res. 2012 Oct;15(6):804–11.

22. Briggs ADM, Wolstenholme J, Blakely T, Scarborough P. Choosing an epidemiological model structure for the economic evaluation of non-communicable disease public health interventions. Popul Health Metr. 2016;14:17.

23. Hennessy DA, Flanagan WM, Tanuseputro P, Bennett C, Tuna M, Kopec J, et al. The Population Health Model (POHEM): an overview of rationale, methods and applications. Popul Health Metr. 2015;13:24.

24. Swanney MP, Ruppel G, Enright PL, Pedersen OF, Crapo RO, Miller MR, et al. Using the lower limit of normal for the FEV1/FVC ratio reduces the misclassification of airway obstruction. Thorax. 2008 Dec;63(12):1046–51.

25. Tan WC, Bourbeau J, Hernandez P, Chapman K, Cowie R, FitzGerald MJ, et al. Canadian prediction equations of spirometric lung function for Caucasian adults 20 to 90 years of age: results from the Canadian Obstructive Lung Disease (COLD) study and the Lung Health Canadian Environment (LHCE) study. Can Respir J J Can Thorac Soc. 2011 Dec;18(6):321–6.

26. Freeman J, Hutchison GB. Prevalence, incidence and duration. Am J Epidemiol. 1980 Nov;112(5):707–23.

27. Bourbeau J, Tan WC, Benedetti A, Aaron SD, Chapman KR, Coxson HO, et al. Canadian Cohort Obstructive Lung Disease (CanCOLD): Fulfilling the Need for Longitudinal Observational Studies in COPD. COPD. 2012 Mar 20;

28. Zafari Z, Sin DD, Postma DS, Löfdahl C-G, Vonk J, Bryan S, et al. Individualized prediction of lung-function decline in chronic obstructive pulmonary disease. CMAJ. 2016 Oct 4;188(14):1004–11.

29. Hankinson JL, Odencrantz JR, Fedan KB. Spirometric reference values from a sample of the general U.S. population. Am J Respir Crit Care Med. 1999 Jan;159(1):179–87.

30. Aaron SD, Fergusson D, Marks GB, Suissa S, Vandemheen KL, Doucette S, et al. Counting, analysing and reporting exacerbations of COPD in randomised controlled trials. Thorax. 2008 Feb;63(2):122–8.

31. Hoogendoorn M, Feenstra TL, Hoogenveen RT, Al M, Mölken MR. Association between lung function and exacerbation frequency in patients with COPD. Int J Chron Obstruct Pulmon Dis. 2010 Dec 9;5:435–44.

32. Albert RK, Connett J, Bailey WC, Casaburi R, Cooper JAD, Criner GJ, et al. Azithromycin for prevention of exacerbations of COPD. N Engl J Med. 2011 Aug 25;365(8):689–98.

33. Hurst JR, Vestbo J, Anzueto A, Locantore N, Müllerova H, Tal-Singer R, et al. Susceptibility to exacerbation in chronic obstructive pulmonary disease. N Engl J Med. 2010 Sep 16;363(12):1128–38.

34. Sadatsafavi M, Sin DD, Zafari Z, Criner G, Connett JE, Lazarus S, et al. The Association Between Rate and Severity of Exacerbations in Chronic Obstructive Pulmonary Disease: An Application of a Joint Frailty-Logistic Model. Am J Epidemiol. 2016 Nov 1;184(9):681–9.

35. Statistics canada. Life tables, Canada and provinces and territories [Internet]. Available from: http://www.statcan.gc.ca/pub/84-537-x/4064441-eng.htm

36. Spencer M, Briggs A, Grossman R, Rance L. Development of an economic model to assess the cost effectiveness of treatment interventions for chronic obstructive pulmonary disease. PharmacoEconomics. 2005;23(6):619–37.

37. Mittmann N, Kuramoto L, Seung SJ, Haddon JM, Bradley-Kennedy C, FitzGerald JM. The cost of moderate and severe COPD exacerbations to the Canadian healthcare system. Respir Med. 2008 Mar;102(3):413–21.

38. Statistics Canada. Modgen (Model generator) [Internet]. [cited 2012 Mar 19]. Available from: http://www.statcan.gc.ca/microsimulation/modgen/modgen-eng.htm

39. R Core Team. R: A Language and Environment for Statistical Computing [Internet]. Vienna, Austria: R Foundation for Statistical Computing; 2017. Available from: https://www.R-project.org/

40. Eddy DM, Hollingworth W, Caro JJ, Tsevat J, McDonald KM, Wong JB, et al. Model transparency and validation: a report of the ISPOR-SMDM Modeling Good Research Practices Task Force--7. Value Health J Int Soc Pharmacoeconomics Outcomes Res. 2012 Oct;15(6):843–50.

41. Calverley PMA, Anderson JA, Celli B, Ferguson GT, Jenkins C, Jones PW, et al. Salmeterol and fluticasone propionate and survival in chronic obstructive pulmonary disease. N Engl J Med. 2007 Feb 22;356(8):775–89.

42. Tashkin DP, Celli B, Senn S, Burkhart D, Kesten S, Menjoge S, et al. A 4-year trial of tiotropium in chronic obstructive pulmonary disease. N Engl J Med. 2008 Oct 9;359(15):1543–54.

43. Asukai Y, Baldwin M, Fonseca T, Gray A, Mungapen L, Price D. Improving clinical reality in chronic obstructive pulmonary disease economic modelling: development and validation of a micro-simulation approach. PharmacoEconomics. 2013 Feb;31(2):151–61.

44. Borg S, Ericsson A, Wedzicha J, Gulsvik A, Lundbäck B, Donaldson GC, et al. A computer simulation model of the natural history and economic impact of chronic obstructive pulmonary disease. Value Health J Int Soc Pharmacoeconomics Outcomes Res. 2004 Apr;7(2):153–67.

45. Briggs AH, Baker T, Risebrough NA, Chambers M, Gonzalez-McQuire S, Ismaila AS, et al. Development of the Galaxy Chronic Obstructive Pulmonary Disease (COPD) Model Using Data from ECLIPSE: Internal Validation of a Linked-Equations Cohort Model. Med Decis Mak Int J Soc Med Decis Mak. 2017 May;37(4):469–80.

46. Menn P, Leidl R, Holle R. A lifetime Markov model for the economic evaluation of chronic obstructive pulmonary disease. PharmacoEconomics. 2012 Sep 1;30(9):825–40.

47. Samyshkin Y, Schlunegger M, Haefliger S, Ledderhose S, Radford M. Cost-effectiveness of roflumilast in combination with bronchodilator therapies in patients with severe and very severe COPD in Switzerland. Int J Chron Obstruct Pulmon Dis. 2013;8:79–87.

48. Menzies NA, Soeteman DI, Pandya A, Kim JJ. Bayesian Methods for Calibrating Health Policy Models: A Tutorial. PharmacoEconomics. 2017 Jun;35(6):613–24.

49. Zafari Z, Bryan S, Sin DD, Conte T, Khakban R, Sadatsafavi M. A Systematic Review of Health Economics Simulation Models of Chronic Obstructive Pulmonary Disease. Value Health J Int Soc Pharmacoeconomics Outcomes Res. 2017 Jan;20(1):152–62.

50. Hoogendoorn M, Feenstra TL, Asukai Y, Briggs AH, Borg S, Dal Negro RW, et al. Patient Heterogeneity in Health Economic Decision Models for Chronic Obstructive Pulmonary Disease: Are Current Models Suitable to Evaluate Personalized Medicine? Value Health J Int Soc Pharmacoeconomics Outcomes Res. 2016 Oct;19(6):800–10.

51. Hoogendoorn M, Feenstra TL, Asukai Y, Briggs AH, Hansen RN, Leidl R, et al. External Validation of Health Economic Decision Models for Chronic Obstructive Pulmonary Disease (COPD): Report of the Third COPD Modeling Meeting. Value Health J Int Soc Pharmacoeconomics Outcomes Res. 2017 Mar;20(3):397–403.

52. Vestbo J, Anderson W, Coxson HO, Crim C, Dawber F, Edwards L, et al. Evaluation of COPD Longitudinally to Identify Predictive Surrogate End-points (ECLIPSE). Eur Respir J. 2008 Apr;31(4):869–73.

53. Tabberer M, Gonzalez-McQuire S, Muellerova H, Briggs AH, Rutten-van Mölken MPMH, Chambers M, et al. Development of a Conceptual Model of Disease Progression for Use in Economic Modeling of Chronic Obstructive Pulmonary Disease. Med Decis Mak Int J Soc Med Decis Mak. 2017;37(4):440–52.

54. Flattet Y, Garin N, Serratrice J, Perrier A, Stirnemann J, Carballo S. Determining prognosis in acute exacerbation of COPD. Int J Chron Obstruct Pulmon Dis. 2017;12:467–75.

55. Chapman KR, Bourbeau J, Rance L. The burden of COPD in Canada: results from the Confronting COPD survey. Respir Med. 2003 Mar;97 Suppl C:S23–31.

56. Canadian Institute of Health Information (CIHI). Care in Canadian ICUs [Internet]. [cited 2018 Oct 26]. Available from: https://secure.cihi.ca/free_products/ICU_Report_EN.pdf

57. Rutten-van Mölken MPMH, Hoogendoorn M, Lamers LM. Holistic preferences for 1-year health profiles describing fluctuations in health: the case of chronic obstructive pulmonary disease. PharmacoEconomics. 2009;27(6):465–77.

